# Atlastin-1 regulates endosomal tubulation and lysosomal proteolysis in human cortical neurons

**DOI:** 10.1101/2024.02.29.582512

**Authors:** Eliska Zlamalova, Catherine Rodger, Francesca Greco, Julia Kleniuk, Aishwarya G. Nadadhur, Zuzana Kadlecova, Evan Reid

## Abstract

Mutation of the *ATL1* gene is one of the most common causes of hereditary spastic paraplegia (HSP), a group of genetic neurodegenerative conditions characterised by distal axonal degeneration of the corticospinal tract axons. Atlastin-1, the protein encoded by *ATL1*, is one of three mammalian atlastins, which are homologous dynamin-like GTPases that control endoplasmic reticulum (ER) morphology by fusing tubules to form the three-way junctions that characterise ER networks. However, it is not clear whether atlastin-1 is required for correct ER morphology in human neurons and if so what the functional consequences of lack of atlastin-1 are. Using CRISPR-inhibition we generated human cortical neurons lacking atlastin-1. We demonstrate that ER morphology was altered in these neurons, with a reduced number of three-way junctions. Neurons lacking atlastin-1 had longer endosomal tubules, suggestive of defective tubule fission. This was accompanied by reduced lysosomal proteolytic capacity. As well as demonstrating that atlastin-1 is required for correct ER morphology in human neurons, our results indicate that lack of a classical ER-shaping protein such as atlastin-1 may cause altered endosomal tubulation and lysosomal proteolytic dysfunction. Furthermore, they strengthen the idea that defective lysosome function contributes to the pathogenesis of a broad group of HSPs, including those where the primary localisation of the protein involved is not at the endolysosomal system.

## Introduction

Hereditary spastic paraplegias (HSPs) are neurological conditions in which the principal clinical feature is progressive spastic paralysis of the legs.[1–3] Pathologically these conditions are associated with a “dying back” axonopathy, in which the longest axons of the corticospinal tract are affected.[4] HSPs are conventionally divided into “pure” and “complex” forms based on the absence or presence of other neurological or non-neurological features.[1] They are single gene disorders and more than 80 HSP loci have been identified so far.[5] Study of the encoded proteins has facilitated mechanistic understanding of the cellular processes involved in HSP pathogenesis and in the normal maintenance of healthy axons. Through this work, HSP proteins have been subdivided into a relatively small number of groups based on their cellular functions, with important subgroups being involved in shaping the endoplasmic reticulum (ER) and in endolysosomal function.[6–8]

The ER is a highly interconnected endomembrane system that consists of sheet-like and polygonal tubular compartments with specialised functions.[9, 10] The anatomy of the tubular ER is governed by sets of shaping proteins (“ER morphogens”) that include the Reticulon, Reep and Atlastin families.[9–15] These proteins are characterised by the presence of two closely spaced membrane domains (often occurring in pairs) followed by an amphipathic helix. The closely spaced membrane domains form an intramembrane “hairpin” structure that is involved in inducing the high membrane curvature typical of ER tubules.[16, 17] The atlastin family of dynamin-like large GTPase proteins have a key function within this ER morphogenic process, as they act enzymatically to homotypically fuse ER tubules to form the network of polygonal structures that characterise the tubular ER.[11, 13, 18] Strikingly, mutations in the genes encoding 4 members of these classes of proteins (reticulon-2, REEP1, REEP2 and atlastin-1) are encoded by genes that are mutated in HSP.[19–22] In addition, these proteins interact at the ER with spastin, a microtubule severing enzyme that is encoded by the gene most frequently mutated in HSP.[12, 23, 24] Spastin is anchored in the ER via a single long hydrophobic sequence that also forms a hairpin membrane domain, and this domain is required for its interaction with other ER shaping proteins.[12] Mutations in the genes encoding spastin, atlastin-1 and REEP1 account for >50% of cases of autosomal dominant pure HSP, the most common subtype in North America and northern Europe. Therefore, understanding how abnormalities of ER morphogenesis affect axonal functions is crucial to our understanding of HSP pathogenesis.[12]

There are three homologous atlastin proteins in mammals, termed atlastins 1-3, which are thought to be at least partially functionally redundant.[25] Atlastin-1 is highly expressed protein in human brain, but less so in other tissues, while atlastins-2 and -3 are minimally expressed in the brain, but highly expressed in many non-neurological tissues.[25] Mutations in atlastin-1 cause a predominantly childhood onset HSP that may be pure but which is often accompanied by peripheral neuropathy - in some families the neuropathy is the most troublesome feature and then the condition is termed autosomal dominant hereditary sensory neuropathy type 1D.[20, 26] The mutational spectrum of atlastin-1-HSP encompasses many missense mutations that tend to cluster at various points within key domains, but also includes a small number of loss of function mutations, including frameshift, nonsense and whole exon deletion mutations.[27] It has generally been thought that most atlastin-1 mutations act via a dominant negative mechanism, although gain of function mutations have been associated with a more severe phenotype and the presence of the loss of function mutations described above suggests that at least in some cases a haploinsufficiency mechanism operates.[27, 28] In this regard, recent *in vivo* data derived from engineering human mutations into *Drosophila* atlastin did not support a dominant negative mechanism and the probability of loss intolerance (pLI) score for the *ATL1* gene is 0.98, indicating that it is intolerant of haploinsufficiency.[29, 30] Furthermore, rare cases of HSP caused by autosomal recessive loss-of-function *ATL1* mutations have been described.[31] *ATL2* has not so far been linked to disease, while mutations in *ATL3* cause a type of autosomal dominant peripheral sensory neuropathy.[32]

Atlastins are essential to generate a proper tubular ER network. Thus non-neuronal mammalian cells lacking atlastins exhibit reduced tubular ER network density accompanied by reduced frequency of ER tubule fusion events. Moreover, an atlastin homologue is one of three components required to reconstitute the tubular ER *in vitro*.[15, 33] The effect of lack or disruption of atlastin on neuronal ER morphology has been studied in invertebrates, which are attractive systems as they have fewer atlastin homologues. Flies lacking atlastin show increased fragmentation of the ER in neurons, including motor nerve terminals.[11, 34] In *C. elegans* neurons lacking functional atlastin, ER sheets expand and rough ER proteins localise ectopically to the axon, while in a GTPase-impaired mutant, the somatodendritic ER network exhibited decreased complexity and ER tubules retracted from higher order dendrites.[35, 36] Surprisingly however, while multiple cellular defects in mammalian neurons lacking atlastin-1 or expressing mutant forms of the protein have been described, and while homozygous knock-in of an atlastin missense mutation caused occasional ER morphology abnormalities in mouse neurons, the effect of lack of atlastin-1 on ER morphology in mammalian, and especially human, neurons has not to our knowledge been reported.[37–39] Thus it is not clear whether atlastin-1 is required for normal ER morphology in mammalian neurons.

Another important group of HSP proteins localises to and is involved in maintaining the correct function of the late endosomal/lysosomal compartments, which are crucial for receptor degradation and autophagic functions.[7, 8] This group includes proteins such as spatacsin (SPG11), spastizin (SPG15) and AP5Z1 (SPG48), which directly function in the endolysosomal system and which are encoded by genes mutated in relatively rare autosomal recessive forms of complex HSP.[40] However, as exemplified by our work on spastin, at least some ER morphogen proteins are also required for normal lysosome function, suggesting that this could contribute more broadly to HSP pathogenesis. Contacts between the ER and endosomes are required for fission of transport tubules that break from endosomes and traffic cargoes (e.g. membrane-associated receptors) to other cellular sites such as the Golgi apparatus.[41–43] We found that spastin is present at these organelle contacts and that defective ER-associated endosomal tubule fission (ETF) in cells lacking spastin caused increased endosomal tubulation accompanied by a block in transport of the mannose 6-phosphate receptor (M6PR) away from endosomes. This receptor normally cycles between the endosomes and the Golgi apparatus, from which it picks up M6P-tagged lysosomal enzymes for delivery back to the endolysosomal system. Consistent with a lack of M6PR at the Golgi, HeLa cells depleted of spastin exhibited lysosomal enzyme mistrafficking, associated with lysosomal enlargement and intralumenal substrate accumulation. We observed very similar lysosomal abnormalities in iPSC-derived neurons from spastin-HSP patients and in neurons from a mouse model of spastin-HSP.[42]

In this study, we set out to answer two questions, i) is atlastin-1 required for normal ER morphology in human neurons? and ii) does deficiency of a classical ER morphogen such as atlastin-1 cause abnormal lysosomal function? We sought to answer these questions in induced pluripotent stem cell (iPSC)-derived human glutamatergic cortical neurons, as these are the target cells of HSP. We found that human neurons lacking atlastin-1 have altered ER morphology, increased endosomal tubulation suggesting an ETF defect and abnormal lysosomal proteolytic activity, supporting the idea that ER and lysosome functions are linked in neurons and that defects in lysosomal function may contribute to the pathogenesis of atlastin-1-HSP.

## Materials and Methods (including Statistical analysis and Data availability) Cell Culture

### Induced pluripotent stem cells (iPSCs)

Human i^3^ iPSCs (generated in a WTC11 iPSC background line) were a kind donation of Michael Ward (NIH, USA). Cells were grown at 37°C and 5% CO2. Specific hood, incubator and other equipment were dedicated to iPSC work. All culturing and iPSC differentiation techniques were as described previously with only minor alterations to the protocol.[44] In short, iPSCs were cultured in TESR-E8 (STEMCELL Technologies) on dishes coated with Matrigel Matrix (Corning). TESR-E8 was replaced daily and cells were passaged at 80% - 90% confluency with 0.5 mM EDTA to maintain colony growth and with the ROCK inhibitor Y-27632 (10 µM, Tocris) to increase cell survival. All cell lines were regularly tested for mycoplasma contamination.

### I^3^Neuron differentiation

Cortical i^3^Neurons were differentiated according to protocols modified from [44]. Briefly, on Day 0 i^3^ iPSCs were dissociated into single cells using StemPro Accutase (Thermo Fisher Scientific) and seeded onto Matrigel-coated plates in Induction Medium (IM) composed of DMEM/F-12 (Thermo Fisher Scientific), 1X N-2 Supplement (Thermo Fisher Scientific), 1X MEM Non-Essential Amino Acids Solution (Thermo Fisher Scientific), 1X GlutaMAX Supplement (Thermo Fisher Scientific), 10 µM Y-27632 (Tocris) and 2 µg/mL doxycycline hydrochloride (Sigma-Aldrich). Pre-differentiated cells were maintained in IM without Y-27632 for 3 days with daily medium changes conducted and NGN2 expression induced using doxycycline. On the third day, cells were replated onto culture plates coated with 0.1 mg/mL poly-L-ornithine (Sigma-Aldrich). From this point, cells were maintained in Cortical Neuron Culture Medium, composed of BrainPhys Neuronal Medium (STEMCELL Technologies), 1X B-27 Supplement (Thermo Fisher Scientific), 10 ng/mL BDNF (PeproTech), 10 ng/mL NT-3 (PeproTech) and 1 µg/mL mouse Laminin (Thermo Fisher Scientific) with half media changes performed every 3-4 days.

## Antibodies

Rabbit anti-SOX2 (2748S, 1:1000), rabbit anti-OCT4 (2750S, 1:1000), rabbit anti-GAPDH (2118, 1:10000), rabbit anti-LAMP1 (D2D11, 1:200) and mouse anti-NANOG (4893S, 1:1000) were obtained from Cell Signalling Technology. Mouse anti-LAMP1 (H4A3, 1:100) was from Santa Cruz Biotechnology, Inc. Rabbit anti-ATL1 (12149-1-AP, 1:500) was from Proteintech group, Inc, rabbit anti-ATL2 (NBP1-78733, 1:1000) was from Novus Biologicals and rabbit anti-ATL3 (A303-313A, 1:2000) was from Bethyl Laboratories. Mouse anti-SNX1 (611482, 1:1000) was from BD transduction laboratories. Mouse anti-MAP2 (ab11268, 1:1000); mouse anti-TAU (ab80579, 1:1000) and rabbit anti-TUJ1 (ab18207, 1:1000) were obtained from Abcam. Rabbit anti-MAP2 (AB5622, 1:100) and rabbit anti-LC3 (L7543, 1:2000) were from Sigma-Aldrich. Rabbit anti–cathepsin D (219361, 1:2000) was from Calbiochem. Mouse anti-Ankyrin G (33-8800, 1:100), goat anti-mouse IgG (H+L) DyLight 800 4X PEG (SA5-35521, 1:7000), goat anti-rabbit IgG (H+L) DyLight 680 (35568, 1:7000) and secondary horseradish peroxidase antibodies anti-rat, anti-rabbit and anti-mouse (1:7000) were from Thermo Fisher Scientific. Alexa Fluor 488-, 568-, and 647-labelled antibodies for immunofluorescence (1:300) were from Molecular Probes.

## CRISPRi

### Cloning of ATL1 targeting sgRNA oligos into a relevant plasmid

CRISPRi sgRNAs were cloned into the pKLV-U6gRNA-EF(BbsI)-PGKpuro2ABFP (pKLV) plasmid (a gift from Kosuke Yusa, Addgene plasmid #50946; RRID: Addgene_50946).[45] The oligo sequences were:

Oligo 1 F: CACCGAACTTCTGCAGCCTGCACG

Oligo 1 R: AAACCGTGCAGGCTGCAGAAGTTC

Oligo 2 F: CACCGGCGCTCGCTGCCTTCTCCG

Oligo 2 R: AAACCGGAGAAGGCAGCGAGCGCC

The pKLV plasmid was digested by the BpiI restriction enzyme (Thermo Fisher Scientific - FD1014). Then, the oligos forming sgRNA duplex were annealed. Next, they were ligated into the open vector by T4 ligase (New England BioLabs). The ligated vector was transformed into competent *E. coli*, grown and minipreped.

### CRISPRi lentivirus production and transduction of iPSCs

For the lentivirus preparation, 80% confluent HEK293T cells growing in antibiotic-free Dulbecco’s modified Eagle’s medium (DMEM) with 4500 mg/l glucose, sodium pyruvate, and sodium bicarbonate (Sigma-Aldrich – D6546) supplemented with 10% fetal bovine serum (FBS, Sigma-Aldrich – F7524) and 2 mM L-glutamine (Sigma-Aldrich – G7513) were transfected with three constructs. The pKLV plasmid containing specific sgRNA oligos, pCMVΔ8.91 and pMD.G plasmids were mixed at a ratio of 50:35:15 to a total of 2 μg of DNA. The DNA was added to 50 μl of Opti-MEM. In another tube, 8 μl of TransIT-293 (Mirus) were mixed with 200 μl Opti-MEM. After a 5 min incubation, both solutions were combined, gently mixed and incubated at room temperature for 20 min. The transfection mix was pipetted dropwise to cells, which were subsequently placed into a category 2 incubator. 24 hours later, the medium on transfected HEK293T cells was exchanged for E8 iPSC media. At 48 hours post-transfection, the viral supernatant was collected, filtered through a 0.45 μm filter, and diluted 1:1 with E8 media. Next, polybrene at a final concentration of 10 μg/ml and Y-27632 ROCK inhibitor (Tocris) at a final concentration of 10 μM were added. Meanwhile, iPSCs were dissociated with Accutase and re-plated as described in [44]. The viral solution was added directly to cells in one well of a 6-well plate. Finally, the iPSCs were span at 1800 rpm for 1 hour at 32°C and then incubated with lentivirus overnight. Transduced cells were selected with puromycin at a concentration of 1 μg/ml for the next two weeks.

## qPCR

QIAGEN RNeasy Mini Kit was used to extract RNA from iPSCs and neurons. High Capacity RNA-to-cDNA Kit (Thermo Fisher Scientific) was employed to transcribe RNA into cDNA following manufacturer protocol. TaqMan qPCR protocol was used in this project. 25 ng of the cDNA (5 ng/μl) were used as a template in each qPCR reaction. The cDNA was mixed with 10 μl of TaqMan Mastermix (Thermo Fisher Scientific - 4304437), 4 μl of H2O and 1 μl of a specific assay containing primers for a gene of interest (Thermo Fisher Scientific). The specific primer sequences are a company trade secret, but all were selected to be exon-spanning probes. Every qPCR assay consisted of a gene expression assay (gene of interest) for each sample, a reference gene control assay (*ACTB* was used) for each sample and non-template control. All reactions were performed in triplicate. The thermocycling conditions were as instructed by the manufacturer. The TaqMan gene expression assays average at 100% amplification efficiency (+/- 10%) which enabled the relative in between-genes comparisons.

## Immunoblotting

Cells were washed twice on ice with PBS and subsequently scraped with ice-cold lysis buffer (1% Triton X-100, 300 mM NaCl, 50 mM Tris [pH 8], 5 mM EDTA [pH 8], protease inhibitors cocktail). The lysis was left to proceed for 15 min on ice. Samples were centrifuged at 20,000 × g for 10 min at 4°C, and the supernatant was collected. The bicinchoninic acid (BCA) assay was performed to determine the protein concentration of each sample. The concentration of samples within one experiment was equalised by the addition of lysis buffer. Cell lysates were kept on ice until further diluted with Laemmli buffer at room temperature. Samples containing Laemmli buffer were boiled for 5 min at 95°C and consequently stored in -70°C to -20°C until western blot analysis. Protein samples were subjected to an SDS–PAGE separation together with SeeBlue Plus2 Prestained Protein Standard (Thermo Fisher Scientific) and transferred onto PVDF membranes. Membranes were blocked in 5% milk with 0.1% Tween in Tris-buffered saline (TBST) for 30-60 min and then incubated in the same solution with primary antibodies at 4°C overnight. Next, membranes were washed three times for 10 min in TBST and incubated with secondary antibodies (in 5% milk in TBST) for approximately 60 min. After subsequent three 10 min washes in TBST, proteins were visualised either with the SuperSignal™ West Pico PLUS Chemiluminescent Substrate (Thermo Fisher Scientific – 34580) for HRP-conjugated antibodies or, for IRDye-conjugated secondary antibodies, on a LI-COR machine with direct infrared fluorescence detection on an Odyssey Infrared Imaging System. Protein levels were quantified using either ImageJ software or LI-COR Odyssey System respectively.

## Cell Viability Assays

### Adenosine triphosphate (ATP) based viability assay

ATP was measured as a proxy of viable cells using the CellTiter-GloR Luminescent Cell Viability Assay (Promega - G7570) according to the manufacturer’s instructions. The kit contains a detergent to lyse viable cells, releasing ATP into the medium. Inhibitors of endogenous enzymes such as ATPase prevent ATP from being degraded. Consequently, with the energy from ATP, a stabilised luciferase uses luciferin to generate luminescence, which can be detected within 10 min using a luminometer. A minimum of 3 wells in a 96-well plate were used as technical repeats for each measurement.

### Lactate dehydrogenase (LDH) based cell dead assay

The CytoTox 96R Non-Radioactive Cytotoxicity Assay (Promega - G1780) was used to measure LDH according to a protocol provided by the manufacturer. LDH is a stable cytosolic enzyme that is released into media upon cell death and lysis. In short, aliquots of the neuron culture media were harvested and subjected to an enzymatic assay during which a tetrazolium salt (iodonitrotetrazolium; INT) is converted into a red formazan product at a rate proportional to LDH concentration.[46] The reaction was stopped after 30 min. The amount of colour was proportional to LDH and in turn to the number of neurons that recently died. The data was collected on a plate reader measuring visible wavelength absorbance at 492 nm. Next, the live neurons, from which media was originally sampled, were lysed. The protocol was repeated with media harvested after all neurons died and released LDH. This measurement correlated with overall cell number and is referred to as the maximum LDH release. Finally, a ratio of LDH spontaneously released into media and the maximum release after complete lysis corresponds to percentage cytotoxicity. A minimum of 3 wells in a 96-well plate were used as technical repeats for each measurement.

## DQ BSA trafficking assay

Neurons were differentiated from iPSCs using the standard protocol. On differentiation day 21, DQ™ Green BSA (Thermo Fisher Scientific - D12050) was added to the cortical media at 10 μg/ml final concentration. It was incubated covered from light for 5 hours in a standard incubator. Then, CellTracker™ Blue CMAC Dye was added into the media at 10 μM final concentration and neurons were incubated for another 20 min. After the second incubation, media was removed, neurons were washed with PBS and fresh media was added to neurons. Immediately after, live imaging was performed in a fully incubated Zeiss LSM 780 confocal laser scanning microscope (37°C, 5% CO2). The control neurons were always imaged second, with the two lines of *ATL1* KD neurons processed first and third. This was to ensure the timing did not bias the result. The acquired images were analysed in ImageJ software. The CellTracker™Blue CMAC Dye stained the entire cells and was used to outline the soma, which was a region of interest. The degree of DQ-BSA degradation was evaluated in a green channel by measuring mean fluorescence intensity per soma.

## Immunofluorescence microscopy on fixed cells

IPSCs and neurons were grown on 10 mm coverslips (Electron Microscopy Sciences - 72290-04), which were previously washed in 1 M HCl heated to 60°C for 8-12 hours, rinsed extensively in sterile water and stored in 100% ethanol. The cells were washed once with PBS before fixation. Fixing was performed with 4% paraformaldehyde (PFA) made in PBS for 20 min at room temperature. Next, the fixative was removed, and cells were washed with PBS three times. Cells were permeabilised for 30 min at room temperature with 0.1% saponin dissolved in PBS. Subsequently, all coverslips were blocked for 1 hour in PBS with 3% bovine serum albumin (BSA). In order to maintain permeabilisation, 0.05% saponin was added to the blocking solution. Next, coverslips were incubated for 1 hour in primary antibodies diluted in a blocking buffer. The coverslips were then washed sequentially three times in blocking buffer and PBS. Secondary antibodies were diluted in a blocking solution and applied to coverslips for 1 hour. Following that, coverslips were washed three times sequentially in blocking buffer, PBS and water. If required for the experiment, 10 μg/ml HCS CellMask Deep Red Stain (Thermo Fisher Scientific - H32721) was applied for 30 min to visualise the entire cell volume. Samples were washed three times sequentially in blocking buffer, PBS and water. Finally, cells were stained with DAPI (Sigma-Aldrich - D9542, 1 μg/ml) for 4 min and once again washed three times sequentially in blocking buffer, PBS and water. Coverslips were mounted in ProLong Gold antifade reagent (Thermo Fisher Scientific – P36934). The mounted samples were dried at room temperature overnight and processed by microscopy imaging. The microscopes used were LSM880 confocal microscope (100× or 63× NA 1.40 oil immersion objective, 37°C), LSM780 confocal microscope (63× NA 1.40 oil immersion objective, 37°C), AxioImager Z2 Motorized Upright Microscope (100× or 63× NA 1.40 oil immersion objective, room temperature, Axiocam 506), and AxioObserver Z1 inverted widefield microscope (63× NA 1.40 oil immersion objective, room temperature, sCMOS ORCA Flash 4 v2). Images were subsequently processed using ZEN software, Huygens Professional software for deconvolution, ImageJ, Adobe Photoshop, and Adobe Illustrator.

## Transduction of mEmerald-KDEL directly into neurons for ER structure visualisation

The pLV-Tre3g-mEmerald-KDEL lentiviral plasmid was a gift from Stefan Marciniak (Cambridge, UK). For the lentivirus preparation, 80% confluent HEK293T cells growing in a T-175 flask in antibiotic-free DMEM (Sigma-Aldrich – D6546) supplemented with 10% fetal bovine serum (FBS, Sigma-Aldrich – F7524) and 2 mM L-glutamine (Sigma-Aldrich – G7513) were transfected with four plasmids. 2.5 μg of pLV-Tre3g-mEmerald-KDEL and each of three packaging plasmids (Addgene - 12251, 12253, 12259) were mixed with 0.5 ml of Opti-MEM. In another tube, 60 μl of PEI (10 mg/ml) were mixed with 440 μl of Opti-MEM. After a 5 min incubation, the PEI-containing solution was dropwise added into the plasmid solution, gently mixed and incubated at room temperature for 30 min. The transfection mix was pipetted dropwise to cells, which were placed into a category 2 incubator. 24 hours later, the medium on transfected HEK293T cells was exchanged. At 72 hours post-transfection, the viral supernatant was collected and filtered through a 0.45 μm filter. Lenti-X Concentrator (Takara - 631232) was used to concentrate the virus. Lenti-X solution was mixed with the lentivirus-containing media at a 1:3 ratio, incubated on ice for 20 min and centrifuged at 4°C for 45 min. The virus pellet was resuspended in 250 μl of PBS, aliquoted and snap-frozen in dry ice before being stored at -70°C. On day 11 of the neuron differentiation, the mEmerald-KDEL lentivirus was directly pipetted into neuron culture media (1 μl per 100000 neurons in an 8-well glass bottom slide, 1.0 cm² per well growth area, Thistle Scientific - IB-80807). The pLV-Tre3g-mEmerald-KDEL is doxycycline-inducible. Therefore, doxycycline-supplemented media (2 μg/ml) was changed daily until day 14 when neurons were imaged live.

## Statistical analysis

Statistical analyses were conducted using Microsoft Excel and GraphPad Prism 7. Averages of independent experiments (each representing separate neuronal differentiation) were compared by statistical tests. The One-way ANOVA (unpaired) or Repeated measures (RM) one-way ANOVA (paired) were employed to compare independent experiment data originating from more than 2 sample conditions. Dunnett’s post hoc test was used to correct for multiple comparisons. The levels of significance were denoted as n.s.-non-significant, p>0.05; *, p<0.05; **, p<0.01; ***, p<0.001; ****, p<0.0001. Graphs presented in the study were generated in GraphPad Prism. Error bars displayed in graphs represent the standard error of the mean (SEM).

## Data availability

Primary data are available by request from the authors.

## Results

### Atlastin-1 becomes the predominant atlastin protein early in human cortical neuron differentiation

We began by characterising the expression of different atlastins in human cortical neurons, the target cells of HSP. To do this we used the integrated, inducible and isogenic (i^3^) iPSC system. In this system, an NGN2 neurogenic transcription factor gene placed under a doxycycline-responsive promoter is knocked in at the AAVS1 safe harbour locus, in the well-characterised stem cell line WTC11.[46] This allows rapid, scalable and reproducible differentiation of i^3^ iPSCs to predominantly glutamatergic cortical neurons (i^3^Ns), upon culture for 3 days in the presence of doxycycline followed by neuronal maturation (Figure 1A). After 14 days these cells express neuron-specific markers and have neuronal morphological characteristics (Figure 1A-C).[47]

**Figure 1.**
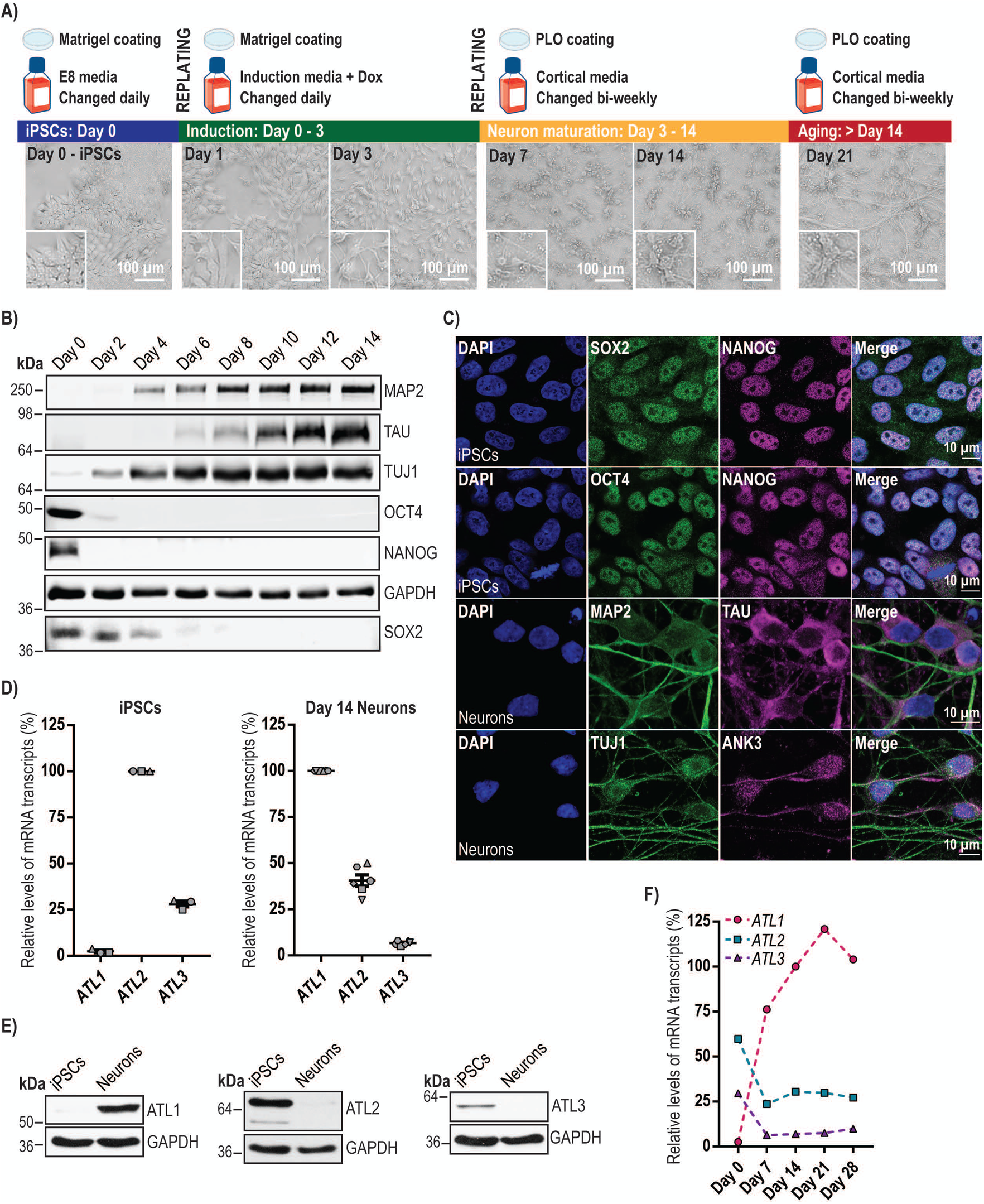
Atlastin expression in human iPSC-derived cortical neurons. **A)** Schematic diagram of the i^3^N differentiation protocol, showing the media and well/coverslip coatings that were used during the protocol. The corresponding brightfield microscope images illustrate cellular morphology during the differentiation period. **B**) i^3^Ns were incubated with doxycycline for 3 days and cultured for the times indicated. Cells were lysed and immunoblotted with the pluripotency (SOX2, NANOG and OCT4) or neuronal differentiation (MAP2, TAU, TUJ1) markers shown. GAPDH blotting serves as a loading control. **C**) iPSCs or day 14 i^3^Ns were fixed, processed for immunofluorescence microscopy and labelled with the pluripotency and neuronal differentiation markers shown. Ankyrin G (ANK3) marks the initial segment of axons (arrows). **D**) Relative mRNA abundance of atlastin gene transcripts in iPSCs or day 14 i^3^Ns, measured by qPCR. Atlastin transcript levels were normalised against the *ACTB* reference gene. Data are displayed relative to the most abundant paralogue in each case. Error bars show mean values +/- SEM of 3 or 6 biological repeats, each carried out in triplicate. **E**) Immunoblots showing the abundance of atlastin proteins in iPSCs or day 14 i^3^Ns. GAPDH blotting serves as a loading control. **F**) The relative mRNA abundance of atlastin transcripts throughout differentiation of i^3^Ns up to 28 days old, measured by qPCR. *ACTB* was used to normalise *ATL1*, -2 and -3 transcript levels. The data points represent percentages relative to the *ATL1* value on day 14. Assays were carried out on one differentiation in triplicate.

We examined the expression of *ATL1-3* transcripts in i^3^ iPSCs and in day 14 (from commencement of induction) i^3^Ns by qPCR. We found that *ATL2* was the predominantly expressed gene in iPSCs; relative to *ATL2* expression, *ATL1* expression was very low (<3%) and *ATL3* expression was substantially less (c.28%). However, in 14 day i^3^Ns *ATL1* mRNA expression had become predominant, with the relative abundance of *ATL2* and *ATL3* mRNA being c.41% and c.7% of that of *ATL1* (Figure 1D). Immunoblotting results reflected these findings, with atlastin-1 being much more strongly expressed in neurons than in iPSCs, while atlastins -2 and -3 were very weakly expressed in day 14 neurons (Figure 1E).

We then carried out a time course experiment to determine the relative abundance of atlastin transcripts throughout a differentiation protocol ending at 28 day old neurons, using qPCR. The switch to *ATL1* becoming the predominant atlastin gene transcript happened by day 7 of differentiation and was maintained at all subsequent time points, while atlastin-2 and -3 expression was reduced from day 7 (Figure 1F). These data are consistent with the previously reported observation that atlastin-1 is the dominant atlastin in human brain and in stem cell-derived human forebrain cortical neurons generated by small molecule-mediated differentiation protocols.[25, 38]

### Generating iPSC-derived human neurons lacking atlastin-1

We next generated i^3^ iPSCs lacking atlastin-1, using a version of the i^3^N system that is knocked-in at a safe harbour locus for CRISPR-inhibition (CRISPRi) machinery. In this CRISPRi system, a dead Cas9 enzyme fused to the KRAB transcriptional repressor domain is constitutively produced in the cells. Lentiviral transduction of a single guide RNA (sgRNA) is used to target the repressor fusion protein to the transcriptional start site of the gene of interest, so durably inhibiting transcription. This typically results in potent depletion of the target protein, with fewer off-target effects than traditional DNA-cutting CRISPR methods or RNAi approaches, and no measurable toxicity from DNA damage.[48] We used this system to make two i^3^ iPSC lines depleted of atlastin-1, each expressing a different targeting sgRNA (lines i^3^N^oligo^ ^1^ and i^3^N^oligo^ ^2^), as well as a control line expressing a scrambled sgRNA (i^3^N^scr^). We then examined the expression of atlastin transcripts and proteins after neuronal differentiation in these lines. QPCR studies confirmed that atlastin-1 mRNA expression in the i^3^N^oligo^ ^1^ and i^3^N^oligo^ ^2^ lines was almost undetectable and <1% of that seen in the i^3^N^scr^ line (Figure 2A). These findings were supported by immunoblotting, which revealed no detectable atlastin-1 in the i^3^N^oligo^ ^1^ or i^3^N^oligo^ ^2^ lines (Figure 2B). In contrast, atlastin-2 and atlastin-3 mRNA (Figure 2C, D) or protein expression (Figure 2E) in iPSCs or i^3^Ns was not affected by atlastin-1 depletion and at the protein level was undetectable in i^3^Ns (Figure 2E). Thus the i^3^N^oligo^ lines were potently depleted for atlastin-1 and had very little atlastin-2 or -3 expression at the protein level.

**Figure 2.**
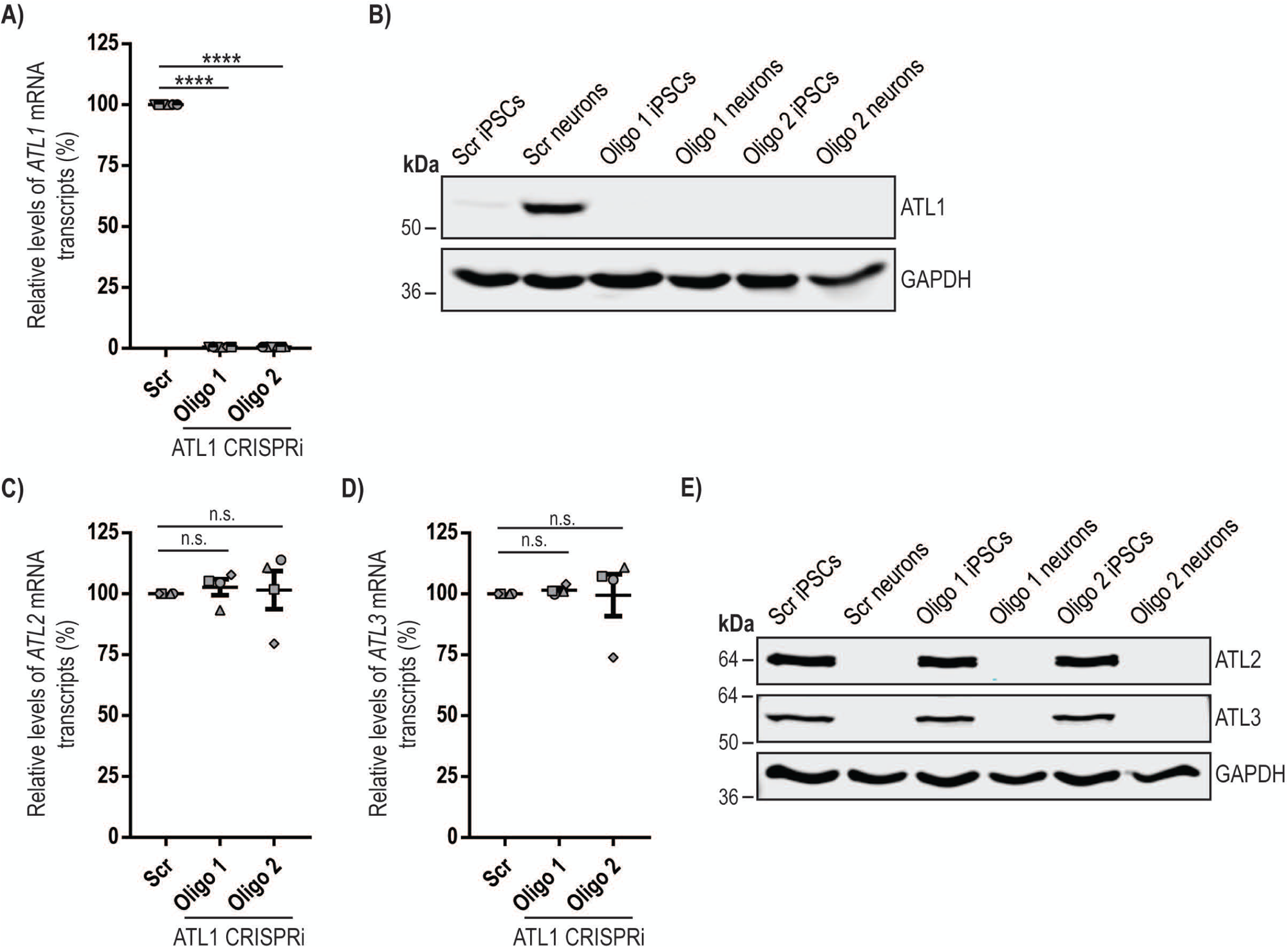
Generation and characterisation of human neurons lacking atlastin-1. **A)** Relative mRNA abundance of *ATL1* transcript in day 14 CRISPRi-i^3^Ns (right) expressing a scrambled sgRNA, or *ATL1*-targeting sgRNAs oligo 1 or oligo 2, measured by qPCR. Data are displayed relative to the scrambled value. Error bars show mean values +/- SEM of 8 i^3^Ns biological repeats, each carried out in triplicate. **B**) Immunoblots showing the abundance of atlastin-1 proteins in iPSCs or day 14 day i^3^Ns expressing the sgRNAs indicated. GAPDH blotting serves as a loading control. **C** and **D**) Relative mRNA abundance of *ATL2* (**C**) or *ATL3* (**D**) transcript in day 14 i^3^Ns expressing the sgRNAs shown, measured by qPCR. Data are displayed relative to the scrambled value. Error bars show mean values +/- SEM of 4 biological repeats, each carried out in triplicate. In all qPCR experiments *ATL* transcript levels were normalised against the *ACTB* reference gene. **E)** Representative immunoblots showing the abundance of atlastin proteins in iPSCs or day 14 i^3^Ns generated from the CRISPRi lines shown. GAPDH blotting serves as a loading control. In A), C) and D) statistical comparisons were carried out with one-way ANOVA with Dunnett’s test for multiple comparisons. n.s., p>0.05; ****, p<0.0001.

### Depletion of atlastin-1 does not affect human neuronal viability

We began to characterise the effects of atlastin-1 depletion in neurons by examining cell viability. We employed two assays, i) the CellTitre-Glo cell viability assay (Promega), which compares the number of live cells by lysing them to release ATP that drives a luciferase reaction, and ii) the lactate dehydrogenase (LDH) viability assay, in which LDH released from dying cells initiates a two-step colourimetric reaction. No difference in day 14 neuron viability between the i^3^N^scr^ line and the two atlastin-1 depleted lines was detected by either assay (Figure 3A, B), so atlastin-1 is not essential for neuronal survival.

**Figure 3.**
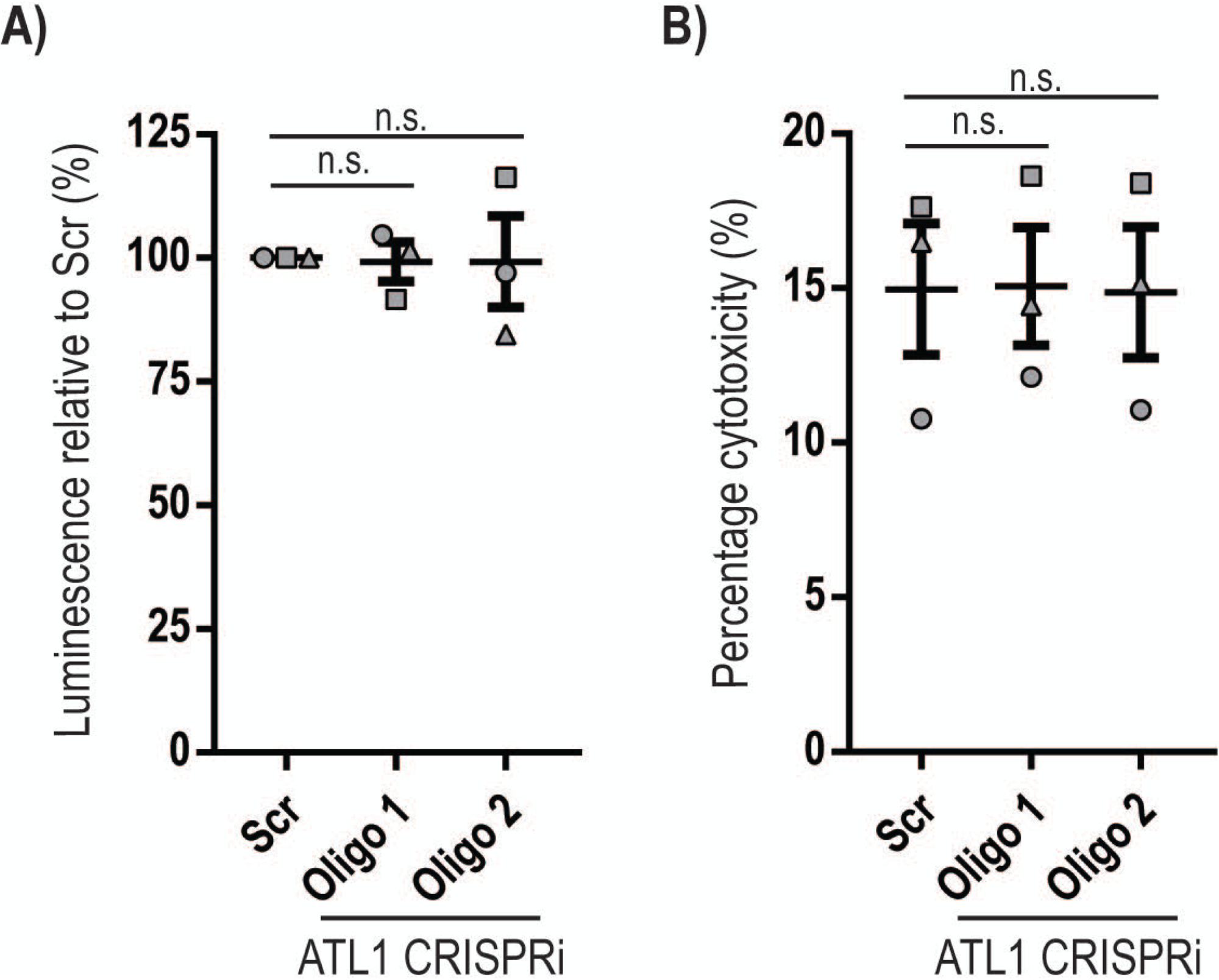
Depletion of atlastin-1 does not affect human neuronal viability. The scrambled and atlastin-1 CRISPRi cell lines indicated were seeded at an equal confluence and differentiated to neurons for 14 days. Cell viability was then quantified using two different assays. **A**) ATP abundance was quantified using the CellTitre-Glo cell viability assay. **B**) LDH levels were quantified in cell medium and after lysis of cultures using the CytoTox 96® Non-Radioactive Cytotoxicity Assay and the ratio of LDH released by spontaneously dying neurons was divided by the total LDH released after complete neuronal culture lysis, to produce a percentage cytotoxicity ratio. In A) and B) results were normalised to the Scr level and mean values +/- SEM are plotted. Statistical comparisons were carried out with one-way ANOVA with Dunnett’s test for multiple comparisons, N=3 biological repeats, each performed with technical triplicates. n.s., p>0.05.

### Neurons lacking atlastin-1 have altered ER morphology in the cell body but not distal neurites

We next analysed ER morphology in the cell body and distal axonal regions of i^3^Ns, using SIM^2^ super-resolution live cell microscopy to examine ER labelled with mEmerald-KDEL (Figure 4A). ER tubules were identified, traced, skeletonised and quantified using ImageJ software (Figure 4B). In the cell bodies, the number of three-way junctions per μ^2^ was modestly but significantly reduced in the atlastin-depleted lines (Figure 4C). However, no significant difference was identified in the ER structure within distal axonal growth cones (Figure 4D, E). The morphological alterations were not associated with any obvious alteration of basal or tunicamycin-activated ER stress pathways (Supplementary Figure 1).

**Figure 4.**
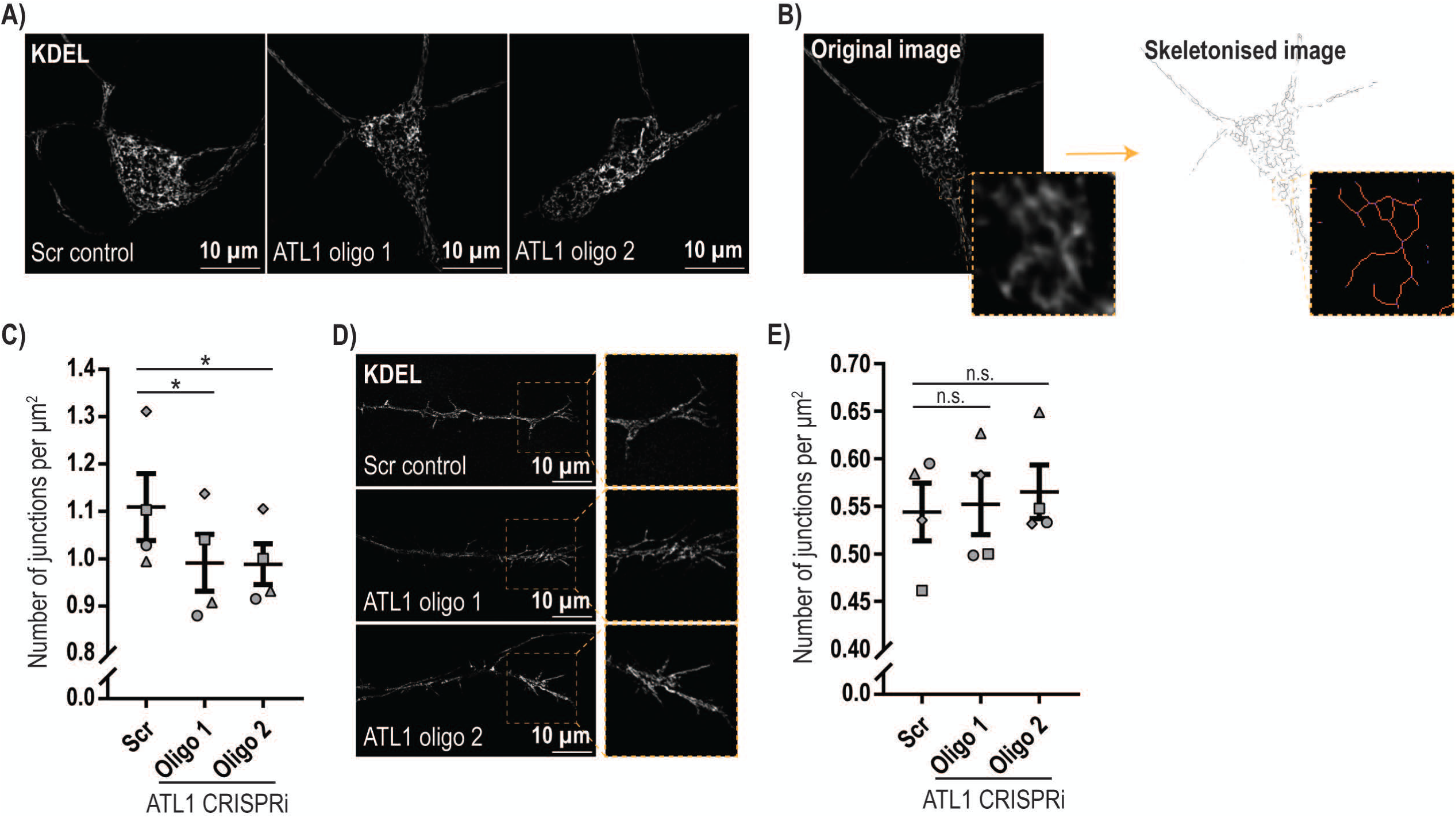
Depletion of atlastin-1 causes reduced ER three-way junction abundance in human neurons. **A)** i^3^Ns from the CRISPRi lines indicated were transduced with a doxycycline-inducible mEmerald-KDEL lentiviral construct and treated with doxycycline for three days from day 11 of differentiation. They were then imaged live with a Zeiss Elyra 7 microscope with Lattice SIM² super-resolution on day 14. Representative images are shown. **B**) ER morphology was analysed using Image J software. Images were analysed blind. Each image was thresholded to select the KDEL signal and then skeletonised to show the network simplified into single pixel wide lines. The zoomed-in skeleton area on the bottom right shows representative skeleton labelling post-analysis. End-point voxels (blue) are pixels with only 1 neighbouring labelled pixel; junction voxels in purple are pixels with more than 2 neighbours; and slab voxels in orange have exactly 2 neighbours. A neighbouring group of junction voxels is counted as one junction. **C**) Junctions were counted in neuronal bodies and the number of junctions was normalised to the cell area. Four neuronal differentiation biological repeats were analysed with at least 25 neurons per genotype in each repeat, with the exception of one repeat for oligo 1 which was based on 4 neurons. Bars show mean +/- SEM. **D**) A similar analysis of ER morphology to that shown in B) was carried out in growth cone regions. The images show representative micrographs of growth cones labelled with mEmerald-KDEL in the i^3^N lines indicated. **E**) Four neuronal differentiation biological repeats were analysed with a minimum of 6 (mean of 27.7) growth cones per genotype analysed in each repeat. The mean number of junctions normalised to the growth cone area was plotted +/- SEM. In C) and E) statistical comparisons were performed with repeated measure one-way ANOVA, with Dunnett’s correction for multiple testing. n.s., p>0.05; *, p<0.05.

In summary, these data show that lack of atlastin-1 alters ER morphology by modestly reducing three-way junction formation in the cell bodies of human neurons.

### Increased endosomal tubulation in neurons lacking atlastin-1

As ETF occurs at ER contact sites, and as atlastin-1’s binding partner spastin promotes ETF, we examined whether the disrupted ER morphology in i^3^Ns was associated with altered endosomal tubulation. Reduced efficiency of ETF leads to a steady state increase in the length of tubules that are attached to endosomes and so we examined whether this phenotype was present by visualising endogenous SNX1, a marker of endosomal tubules that traffic from endosomes to the Golgi apparatus, in the cell bodies of day 14 i^3^Ns (Figure 5A).[49] We found that i^3^N lines lacking atlastin-1 had a significant increase in both the percentage of neurons that had at least one SNX1-positive endosomal tubule >1 μm long and in the length of the longest single endosomal tubule per cell, compared to the i^3^N^scr^ control line (Figure 5B). This was not associated with an increase in SNX1 protein abundance, as measured by immunoblotting (Figure 5C).

**Figure 5.**
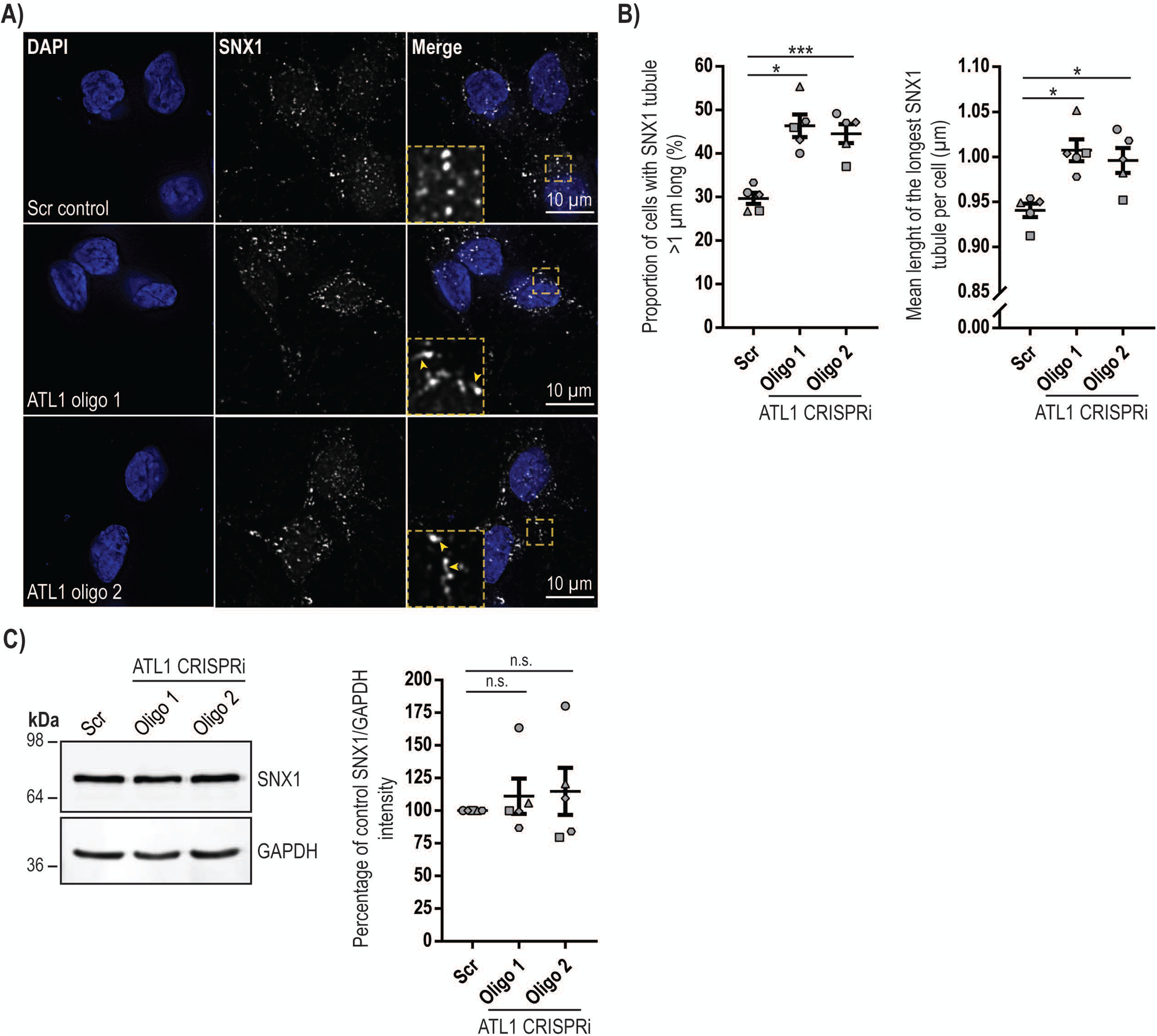
Human neurons lacking atlastin-1 show increased endosomal tubulation. **A)** Fixed day 14 neurons from the CRISPRi-i^3^N lines indicated were immunolabelled for SNX1 and imaged with a Zeiss AxioImager Z2 Motorized Upright Microscope and the images were deconvolved using Huygens software. DAPI was used to visualise nuclei. Representative images of SNX1-labelled endosomes and endosomal tubules are shown. In the right-hand panels the magnified images correspond to the boxed regions in the main images. **B**) The images were blinded for analysis and the longest tubule per cell was measured. The percentage of cells containing a tubule over 1 µm in length is plotted in the left hand chart, while the mean length of the longest tubule per cell is plotted in the right hand chart. The experiment was repeated 5 times with a minimum of 52 neurons analysed per genotype in each differentiation biological repeat. Error bars show SEM. *, p<0.05; ***, p<0.001. **C**) Lysates from day 14 i^3^Ns from the CRISPRi cell lines indicated were immunoblotted for SNX1. GAPDH serves as a control to validate equal lane loading. Immunoblot band intensity was quantified in ImageJ in 5 such experiments and plotted in the corresponding chart. n.s, p>0.05. In B) statistical comparisons were performed with repeated measures one-way ANOVA with Dunnett’s correction for multiple testing. In C) statistical comparisons were performed with one-way ANOVA with Dunnett’s correction for multiple testing.

### Altered lysosomal morphology and proteolytic function in neurons lacking atlastin-1

In non-polarised cells and neurons lacking spastin there is an increase in the size of LAMP1-positive vesicles and disrupted lysosomal function.[39] We therefore examined the percentage of day 14 i^3^Ns that had large LAMP1-positive vesicles (defined as >1.6 μm in diameter) but found no significant difference between the scrambled control and atlastin-depleted lines (Figure 6A, B). We then employed an automated image analysis method to analyse whether the number and average size of vesicles containing the lysosomal enzyme cathepsin D was affected by atlastin-1 depletion. As cathepsin D is contained within vesicles, these parameters are easier to quantify with automated methods than with LAMP1, since the membrane location of LAMP1 means that resolution of individual vesicles is more difficult as signals from different vesicles more commonly overlap. We found a significant increase in the percentage of cells with larger cathepsin-D puncta for the i^3^N^oligo^ ^2^ line, with a trend in the same direction in the i^3^N^oligo^ ^1^ line (Figure 6C). Similarly, there was a significant reduction in the number of cathepsin D puncta per cell in the i^3^N^oligo^ ^2^ line, with a trend in the same direction in the i^3^N^oligo^ ^1^ line (Figure 6D). We concluded that there are subtle alterations in the morphology and number of cathepsin D puncta in i^3^Ns lacking atlastin-1.

**Figure 6.**
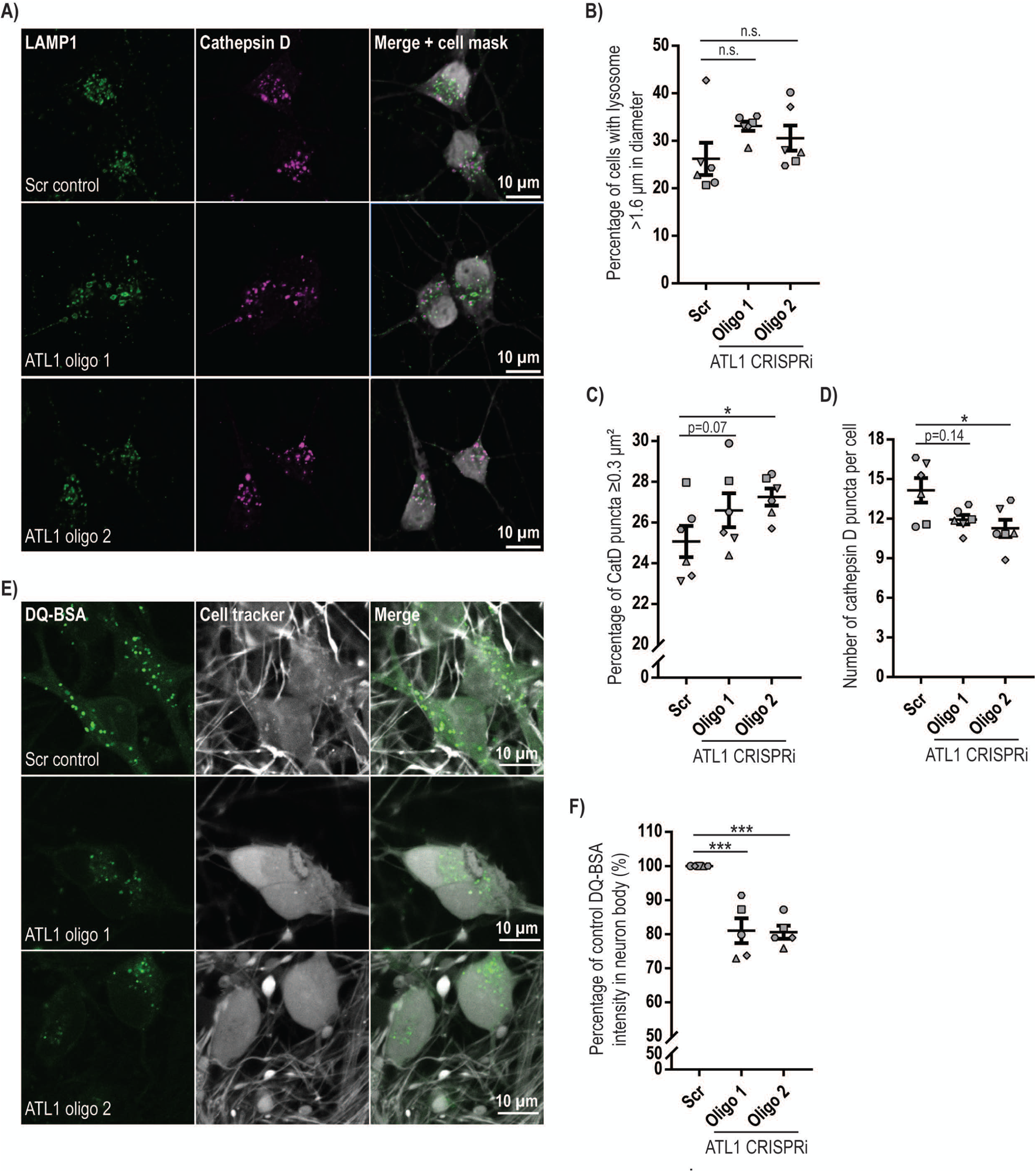
Lysosomal proteolytic abnormalities in cells lacking atlastin-1. **A)** Day 14 i^3^Ns from the lines indicated were fixed and immunolabelled for LAMP1 and cathepsin D, as well as cell mask whole-cell and DAPI nuclear labels. Cells were imaged with a Zeiss Axio Observer microscope with a tile imaging function and the images were deconvolved using Huygens software. **B**) The diameter of the largest LAMP1-positive vesicle per cell was measured in at least 100 cells per genotype and the percentage of neurons with at least one lysosome over 1.6 μm in diameter was calculated and plotted. LAMP1 vesicle size measurement was performed blind to genotype. N=6 biological repeat experiments. **C**) and **D**) The number and area of cathepsin D-positive puncta were measured using an automated protocol developed in Image J software. Only signal within the cell area outlined by the cell mask was considered. Cell number was calculated based on the number of nuclei in the field. The percentage of cells with cathepsin D puncta >0.3 μm^2^ in area is plotted in **C**), while the total number of cathepsin D puncta per neuron is shown in **D**). On average 300 cells were analysed per genotype in each experiment, N=6 biological repeats. Error bars show SEM. n.s., p>0.05; *, p<0.05. **E**) Day 21 neurons from the i^3^N lines indicated were incubated with DQ-BSA substrate for 5.5 hours and with a live-imaging whole cell stain (Cell tracker) for 20 min. Living cells were then imaged with a Zeiss LSM 780 Confocal Microscope. **F**) Neuron somas were selected using the cell tracker channel. The mean DQ-BSA fluorescent intensity was measured and recorded for each cell. Mean results from each experiment are plotted as percentage relative to the Scr control. 5 independent experiments were performed and a minimum of 50 cells per genotype in each experiment was used for quantification. Error bars show mean +/- SEM. ***, p<0.001. In B), C), D) and G) statistical comparisons were made using repeated measures one-way ANOVA with Dunnett’s correction for multiple testing.

In light of these results, we examined lysosomal proteolytic function with the DQ-BSA assay. This assay uses bovine serum albumin that is so heavily labelled with a BODIPY dye that fluorescence is quenched. Upon endocytosis into cells, DQ-BSA is hydrolysed by lysosomal proteolytic activity, relieving quenching and producing a fluorescent signal. We employed this assay in day 21 i^3^Ns, to maximise the chance of detecting any proteolytic dysfunction. We found that 5.5 hours after uptake, DQ-BSA fluorescence intensity was significantly reduced in neurons from both i^3^N^oligo^ lines lacking atlastin-1, at approximately 80% of the control i^3^N^scr^ neuron intensity (Figure 6E, F). This was unlikely to be caused by a defect in endocytosis, as transferrin receptor uptake was not altered by depletion of atlastin-1 (Supplementary Figure 2).

We concluded that atlastin-1 is required for normal lysosomal proteolytic function in neurons. This lysosomal proteolytic defect did not translate to an obvious autophagy defect, as immunoblotting experiments with the autophagy marker LC3 did not identify any consistent abnormality in autophagic flux in neurons lacking atlastin-1 (Supplementary Figure 3).

## Discussion

In this study we examined the role of atlastin-1 in human iPSC-derived i^3^Ns. These neurons have the characteristics of glutamatergic cortical neurons, which reflects the corticospinal tract neurons that are affected in HSP.[47] In i^3^Ns we found that atlastin-1 was the most highly expressed atlastin homologue, in contrast to iPSCs, where atlastin-2 and -3 were both abundant. The switch between these two expression patterns happened within 7 days of NGN2-mediated induction of neuronal differentiation. This atlastin-1 expression pattern is consistent with previous data on human cortical neurons made from patient-derived iPSCs using small molecule-mediated differentiation protocols, where the pattern of strong atlastin-2 and -3 and weak atlastin-1 expression in stem cells was reversed after neuronal differentiation.[38] It is also consistent with tissue expression profiling, which has shown that atlastin-1 is most highly expressed in the brain and rather weakly expressed in other tissues.[25] Thus i^3^Ns correctly model the expected physiological atlastin protein expression pattern in central nervous system neurons.

We used CRISPRi to generate i^3^N lines lacking the protein. We selected a CRISPRi approach as it avoids the DNA damage of cutting CRISPR technologies (which can be toxic to iPSCs), is associated with few off-target effects and often yields very efficient gene silencing. We found that the i^3^N lines we generated had potent atlastin-1 depletion, with >99% suppression of the *ATL1* transcript and no detectable atlastin-1 protein expression. In the atlastin-1 depleted i^3^Ns atlastin-2 and -3 were barely detectable at the protein level, and thus overall these neurons had very low expression of any atlastin proteins.

We observed altered ER morphology in i^3^Ns lacking atlastin-1. To our knowledge, while other studies have demonstrated that expression of mutant atlastin-1 disrupts neuronal ER morphology, this is the first demonstration that lack of atlastin-1 is sufficient to cause an ER morphological defect in mammalian neurons. The ER morphological alteration that we observed was consistent with the known role of atlastins in the homotypic fusion of ER tubules, in that there was a reduction in the number of three-way junctions in the ER of neuronal cell bodies but not in axons. The reduction was subtle, which was somewhat surprising against the background of the low expression of any atlastin protein in i^3^Ns. This suggests that low concentrations of atlastin proteins are sufficient to support homotypic ER tubule fusion, or that atlastin-independent mechanisms for ER tubule fusion exist.

In light of this ER morphology defect, we investigated whether endosomal tubulation was disrupted in i^3^Ns lacking atlastin-1, as abnormalities in ER-associated endosomal tubule fission, coupled to downstream lysosomal abnormalities caused by defective lysosomal enzyme traffic, have been identified in spastin-HSP. Defective endosomal tubule fission results in elongation of endosomal tubules, and we observed an increase in both the proportion of cells with elongated endosomal tubules and the length of the longest tubule in i^3^Ns lacking atlastin-1. Our results underscore the importance of proteins that localise to the ER and regulate ER morphology in governing endosomal tubule events, although it is perhaps surprising that the subtle alteration in ER morphology we saw in atlastin-1-depleted neurons could alone result in the endosomal tubulation we observed. Further studies will be required to understand fully the mechanism of this effect. Nevertheless, atlastin-1 now joins the group of proteins encoded by HSP genes that have been implicated in regulating early endosomal tubulation, which includes spastin and WASH5C.[43, 50]

The disruption of endosomal tubulation was accompanied by findings that indicate disrupted lysosomal enzyme distribution and function in the i^3^Ns lacking atlastin-1, including a reduced number and increased size of cathepsin D-positive puncta, as well as a reduction in the lysosomal proteolytic capacity of the cells as measured by the DQ-BSA assay. A similar reduction in lysosome number has been reported in the muscle of flies lacking atlastin.[51] While further work will be required to understand the mechanism of this effect, we suggest that in neurons lacking atlastin, abnormal endosomal tubule fission affects lysosomal function by disrupting traffic of lysosomal enzymes. Lysosomal enzyme receptors are normally recycled back to the Golgi via endosomal tubules and failure of this process means that they are not available at the Golgi to bind newly synthesised lysosomal enzymes for delivery back to the lysosomal system; lysosomal enzymes are instead secreted.[42] Consistent with this, *Dictyostelium discoideum* amoebae lacking the single atlastin homologue *sey1* showed increased secretion of lysosomal enzymes.[52] Of note, the lysosomal function that we observed was not associated with disruption of generalised autophagy, although this does not exclude participation of atlastin-1 in selective autophagy pathways.

In summary, our results demonstrate novel roles for atlastin-1 in endosomal tubulation and lysosomal function. Thus atlastin-1-HSP joins the group of HSPs in which neuronal lysosomal dysfunction has been identified, and more broadly joins the group of neurodegenerative conditions, including Alzheimer’s disease and Parkinson’s disease, in which lysosomal dysfunction likely plays a pathological role.[53]

## Acknowledgements

We thank Michael Ward for the wild-type i^3^N and i^3^N-CRISPRi lines and for helpful discussions. This research was supported by the CIMR Microscopy Core Facility. We thank Stefan Marciniak for the pLV-Tre3g-mEmerald-KDEL lentiviral plasmid.

