## Supplementary methods and figures for "Atlastin-1 regulates endosomal tubulation and lysosomal proteolysis in human cortical neurons"

##### Transferrin Receptor Endocytosis

Neurons were differentiated and grown in a 96-well stripwell plate (Corning – 9102, $10^5$ neurons per well). This allowed for every plate strip to be handled separately. Endocytic capacity was tested by uptake of anti-transferrin receptor (TFR1) antibody (Bio X Cell - BE0023, 7 µg/ml in neuron cortical medium). The uptake was performed at 37°C for 1, 5, 10, 20, and 30 min. Immediately after, the neurons were washed 13× with ice-cold sodium acetate buffer (0.2 M, pH 5.0) and 3× with ice-cold neutralising buffer (DMEM/F12 - Invitrogen with 0.4% BSA). The neurons were fixed for 25 min (4% PFA, 0.25% glutaraldehyde). Besides the uptake time points, two controls were included in the experiment. First, the total surface TFR1 was measured by neurons being cooled on ice for 15 min to inhibit endocytosis. Then, they were incubated with the anti-TFR1 antibody on ice for 15 min. This was followed by 3 washes with the ice-cold neutralising buffer and fixation. Second, acid wash efficiency was measured by neurons again being cooled on ice for 15 min and incubated with the anti-TFR1 antibody on ice for 15 min. This time, they were washed 13× with the ice-cold sodium acetate buffer and 3× with ice-cold neutralising buffer. This treatment should theoretically only allow surface antibody binding (no endocytosis) and then all the antibodies should be dissociated by the acidic washes. Neurons were fixed as described above.

All fixed neurons were washed 4× post-fixation with the neutralising buffer. They were permeabilised (0.1% saponin in the neutralising buffer, 15-30 min) and incubated with secondary goat anti-mouse HRP antibody (Thermo Fisher Scientific - 41-112-020317, 1:1000 in neutralising buffer with 0.05% saponin, 60 min at room temperature or overnight at 4°C). Then, staining with 10 µg/ml HCS CellMask Deep Red Stain (Thermo Fisher Scientific - H32721, 30 min) was performed. Cells were washed 6× with PBS. To finalise the experiment, the HRP signal was measured. O-phenylenediamine dihydrochloride (OPD; HRP substrate) was used to produce a signal corresponding to the amount of secondary antibody (Sigma-Aldrich - P8287-100TAB, 1 tablet with 10 ml of 0.15 M, pH 5.0 citric buffer and 8 µl of $H_2O_2$). The oxidation of OPD with $H_2O_2$ catalysed by HRP produces a yellow-orange product detectable at 492 nm with a plate reader. The cell number was measured with the plate reader using the CellMask fluorescent signal (680 nm) and used to normalise the HRPproduced signal. A minimum of 3 wells was analysed for each condition as technical repeats, from which results were averaged.

**Autophagy flux**

Differentiating neurons were grown in nutrient-rich cortical media. They were either untreated or exposed to 100 nM Bafilomycin A1 for 24 hours, before being lysed and immunoblotted for LC3-II.

#### Supplementary Figure Legends

**Supplementary Figure 1. Assays of ER stress in human neurons lacking atlastin-1.**
Transcript analysis was carried out by qPCR for the ER stress pathways target genes shown, in the presence and absence of the ER stress inducer tunicamycin, on day 14 CRISPRi $I^3N$ lines. Assays were carried out in technical triplicates on 4 neuronal differentiation biological repeats. Means +/- SEM are plotted. Statistical comparisons were performed with one-way ANOVA, with Dunnett's correction for multiple testing.

**Supplementary Figure 2. Measurement of endocytosis in human neurons lacking atlastin-1.**
**A**) Scheme showing a simplified workflow developed to measure the kinetics of endocytosis in $i^3$ neurons. TFR1 endocytosis was followed in this assay using the uptake of antibodies binding to the TFR1 at the cell surface. After each endocytosis time point, surface antibodies were removed with an acid buffer wash. Hence, only internalised antibodies contribute to the result. Quantification was achieved using HRP-fused secondary antibodies. HRP catalyses the oxidation of OPD into 2,3-diaminophenazine, which is detectable by absorbance measurement at 492 nm. The absorbance is proportional to the internalised TFR1. Furthermore, neurons were stained with a fluorescent whole-cell stain, the intensity of which was measured per well to approximate neuron density and to normalise the results. Two control experiments were performed on ice to inhibit endocytosis; firstly, the total TFR1 available on the cell surface was measured by allowing antibody binding and not removing it by the acid washes and secondly, the same protocol was repeated with the inclusion of the acid washes to measure residual absorbance when no endocytosis takes place and after antibodies are removed from the cell surface. The scheme was created with BioRender.com. **B**) The endocytic assay was performed simultaneously for Scr and atlastin-1 depleted neurons. The primary antibodies were incubated with the neurons at 37$^o$C for 1, 5, 10, 20 and 30 minutes. The background absorbance detected from the acid wash control was subtracted from all results. The absorbance measurements from each time point were also normalised to the total TFR1 available at the cell surface and to the cell number approximated by the cell mask fluorescence. Four independent experiments were

completed and their mean is shown in the plot. Each experiment was performed with 3 technical repeats. Error bars
represent the SEM.

**Supplementary Figure 3. Autophagic flux in human neurons lacking atlastin-1.**

**A**) Day 14 $i^3$Ns were lysed after no treatment or after being treated with 100 nM BafilomycinA1 for 24 hours, then immunoblotted for LC3. GAPDH serves as a loading control. **B**) The plot shows the quantification of the LC3-II intensity normalised to the GAPDH level in the same sample. The results are expressed as percentages relative to the Scr control under basal conditions. 7 independent experiments were analysed. Error bars represent the SEM. Statistical testing was done with one-way ANOVA versus the corresponding Scr condition, n.s., $p>0.05$.

### Supplementary Figure 1

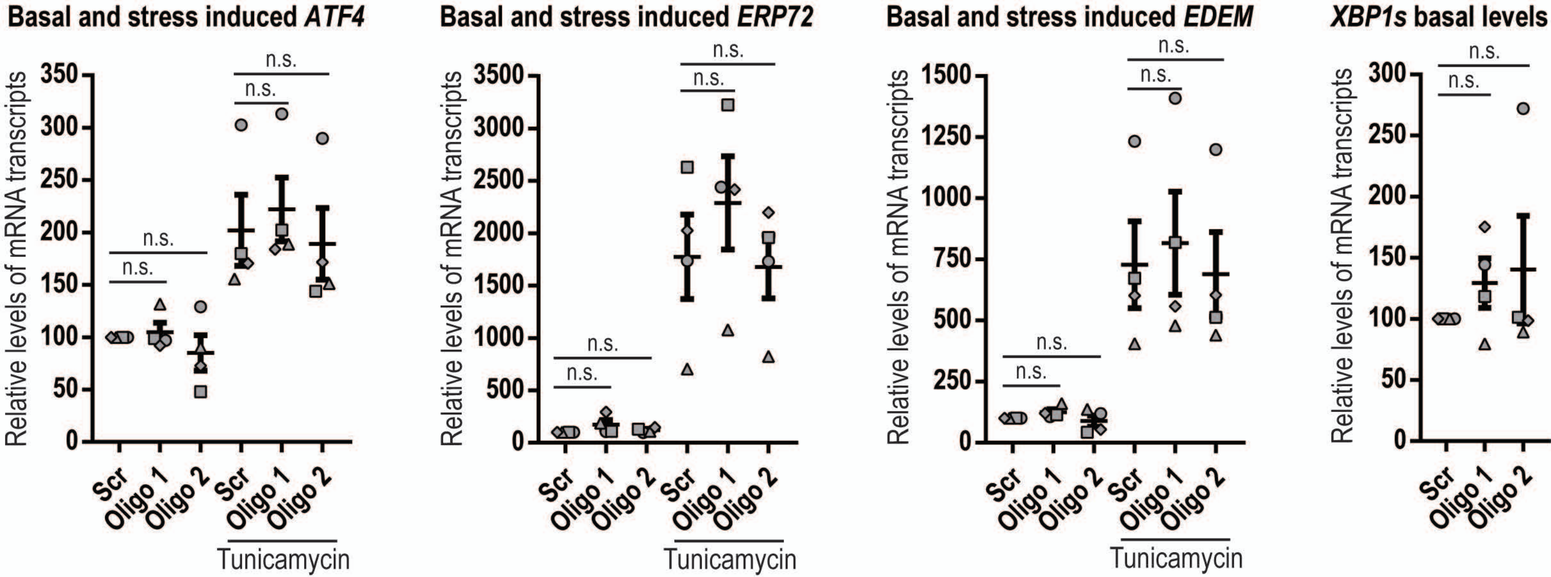

### Supplementary Figure 2

A)

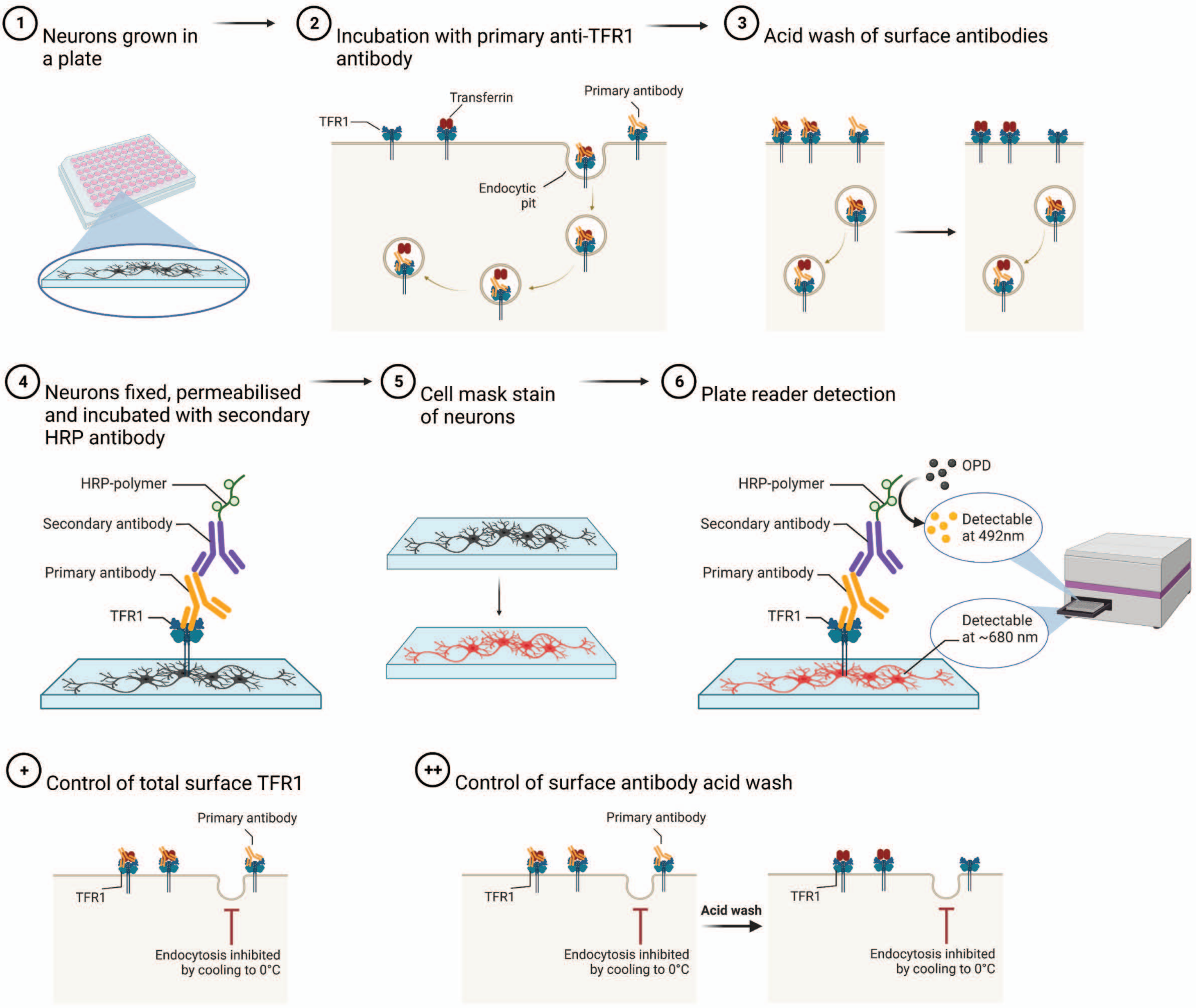


B)

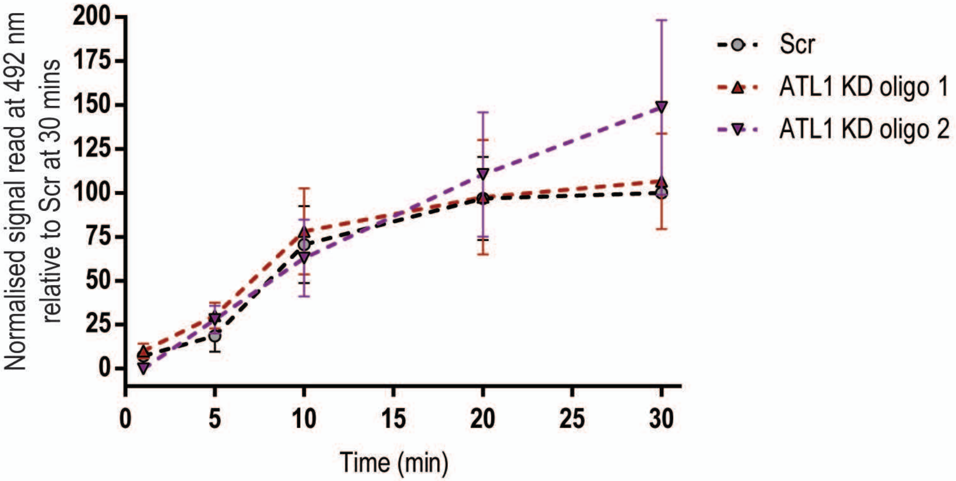

### Supplementary Figure 3

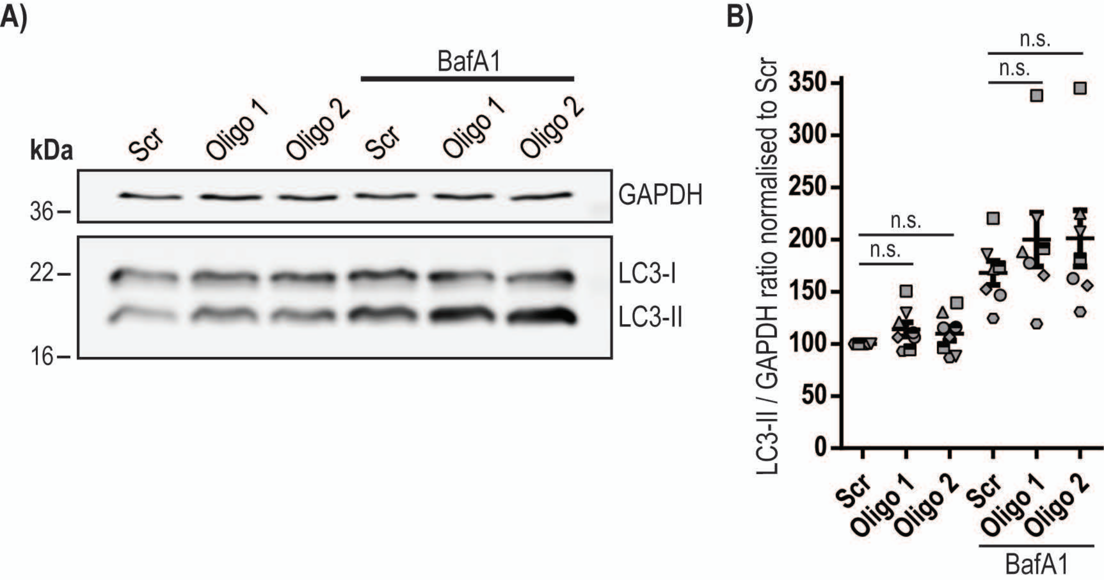
